# Developing an approach to list bryophytes of conservation interests for England’s Local Nature Recovery Strategies

**DOI:** 10.1101/2025.08.01.668143

**Authors:** Michael Davies, Jamie Bojko, Diana Feliciano, Ambroise Baker

## Abstract

Biodiversity is globally declining, and the United Kingdom (UK) recognised as one of the most nature-depleted countries. With an aim to restore ecosystems, the UK has initiated a policy of Local Nature Recovery Strategies (LNRS), to be published for 48 areas across England. We devised and applied a workflow using open access data to produce preliminary lists of bryophytes of conservation interest for each LNRS area. Our objectives include: (i) to ascertain which LNRS area match the current recording scheme based on Watsonian vice-counties; (ii) to produce data-driven draft lists for each LNRS area, (iii) to compare this list in the Tees Valley area with a species list derived from expert knowledge. Using an inclusion threshold at 90% of surface area, we found that 20 out of the 48 LNRS areas are compatible with the vice-county-based scheme. We determine that LNRS areas included on average 9.3 (+/- 6.9) species threatened by extinction, a supplementary 3.6 (+/- 3.4) species near threatened and, adding nationally rare or scarce species, a total of 66.3 (+/- 35.1) species. A case study based on the Tees Valley revealed an overall good match between data and expert driven species lists, despite discrepancies relating to geographical attributions near boundaries and additional species solely selected by expert opinion. It is anticipated that the preliminary lists produced could save valuable time to local experts when working towards LNRS bryophyte lists, and that the approach developed will be relevant to other organism groups, supporting effectively LNRSs across England.

## Introduction

Recognised as one of the most nature depleted nations in the world, the UK has experienced a species abundance decrease by approximately 40%, with average species populations decreasing by 13% since 1970, and 15% of species at risk of extinction (Hayhow et al. 2019; Scott et al. 2024). Species decline is often associated with land-use change, often leading to habitat degradation, fragmentation, and loss (Boatman et al. 2007; Burns et al. 2016). Of the United Kingdom’s (UK’s) terrestrial habitat, 80% is intensively managed for agriculture and urban development, leaving little of natural and semi-natural habitats remaining (Powell and Wentworth 2021). Attempts to reverse biodiversity losses through UK parliamentary action began with the 2010 Lawton review (Lawton et al. 2010), which presented a vision of protection for existing natural habitats, and ensuring that those, alongside restored areas, would follow four habitat conservation principles: bigger, better, more and joined-up (BBMJ). To implement the Lawton Review recommendations at a local level in England, the Environment Act 2021 presented a delivery mechanism named Local Nature Recovery Strategies (UK Government 2021), along with supporting mechanisms beyond the scope of this article. Unlike prior biodiversity schemes centralised at national scale, LNRS are the responsibility of 48 decentralised authorities; a shift designed to improve alignment with other local land-use planning powers, boost utilisation of local expertise, and allow for broader community engagement (Traill-Thompson 2021). Three main results are expected from each LNRS: firstly, agreements around nature recovery and species priorities, secondly mapping of existing local habitats valuable to nature alongside statements providing detail of the regional biodiversity, and thirdly developing maps of habitat improvement and wider environmental proposals (DEFRA 2023). Following the guidance regarding the selection of species priorities for LNRS, lists are to prioritise native species assessed as extinct, regionally extinct, threatened, and near threatened against International Union for the Conservation of Nature (IUCN) criteria, and may contain species suitable for conservation translocation; with stakeholders and public providing supplementary species of conservation importance (Natural England 2023). The IUCN categorise species depending on their extinction risk determined by four criteria: populations declining or projected to do so (Criterion A); narrow geographic distribution and are experiencing fragmented and/or strongly fluctuating or declining populations (Criterion B); small populations in decline (Criterion C); very small or restricted populations (Criterion D) (IUCN 2013). This methodology provides a well-established, widely applied objective assessment tool (Mace 2008). Using IUCN criteria and including input from local stakeholders – from nature groups, to landowners, and the public – matches some suggestions recognised by Crowther et al. (2023) to produce effective nature recovery strategies at local scales and to inform future national biodiversity, nature conservation, and agricultural-environmental schemes.

Bryophytes, the second largest plant group consisting of mosses, liverworts, and hornworts, are understood to be the link between vascular plants and ancestral algae, representing the closest modern relatives to terrestrial pioneer plants (Patino and Vanderpoorten 2018). Despite this, non-vascular plants typically receive less attention than their vascular counterparts, often being overlooked even when they are the dominant group in an environment (Boch et al. 2013; Ye et al. 2023). Noted reasons include a general lack of interest and/or knowledge, difficulty to spot and identify, and a poor understanding of their life traits and ecosystem functions from people with expertise (Eldridge et al. 2023; Slate et al. 2024). Via their evolution journey, bryophytes adapted to surviving in all environments with poikilohydry - absorbing and loosing water through the entire gametophyte (van Zuijlen et al. 2023). Although some species are capable of intaking water from substrates, not relying on root systems for uptake allows for the colonisation of rock faces, tree bark, and other substrates mostly inhospitable to vascular plants (Sokolowska et al. 2017; Patino and Vanderpoorten 2018). The unique water absorption and retention methods of bryophytes act to protect soils and reduce water loss through evaporation, actions that suggest they could be used as pioneer species for ecological restoration efforts (Ye et al. 2023). Consideration in conservation programmes is therefore important to maintain biodiversity of species and function, as well as to measure changes, particularly in forested areas where bryophytes can contribute greatly to plant biomass and species richness (Sun et al. 2015).

The British Bryological Society (BBS) has promoted bryophyte knowledge for over 100 years, with long-lasting focus on taxonomy and biogeography, resulting in Britain and Ireland having one of the best-studied bryophyte flora in the world. Most recent publications are underpinned by species distribution knowledge based on up-to-date taxonomy (Blockeel et al. 2020; Hodgetts et al. 2020), decades of field exploration (Blockeel et al. 2014; Picklington et al. 2023) and a thorough assessment of extinction threat based on IUCN criteria for each known British species (Callaghan 2023).

Based on the BBS rich knowledge of bryophytes diversity in England, this study provides an accessible, reproducible, transferrable methodology capable of generating bryophyte species lists relevant for each of the forty-eight LNRS areas in England. The lists contain species of conservation interest present in each area, their conservation status and whether occurrences are current or historical. By providing a detailed reference point for each LNRS, it is hoped that bryophytes are more likely to be considered in LNRS. The objectives are to:

(1) List bryophyte species of conservation interest occurring in England;
(2) Ascertain which LNRS area match the current recording schemes based on Watsonian vice-counties;
(3) Produce data-driven species lists, using the vice-county-based distributional Census Catalogue (CC) published by the British Bryological Society, when relevant, or otherwise the geographical coordinate-based database curated by the British Bryological Society;
(4) Compare these results for one LNRS area (the Tees Valley) where the bryophyte list had been built using resource-intensive expert knowledge (expert-driven approach).

## Methodology

### Project workflow

The workflow developed for this project is illustrated in Figure 1. To tailor bryophyte species lists for each LNRS areas, we relied on two databases expertly curated by the British Bryological Society (BBS). The BBS maintains lists of species attested for each of the Watsonian vice-counties, most recently updated and published as an interim Census Catalogue (CC) (Pilkington et al. 2023). Vice-counties mapping is based on boundaries first published in 1852 that divide England into 57 vice-counties (Dandy 1969), and remains extensively used in natural history in the UK. However, LNRS boundaries may not systematically match those of the vice-counties and in this case the CC may not be fully relevant. For LNRS area where the CC was less relevant, we relied on a second BBS database, recording bryophyte species occurrence using geographical coordinates, in particular the Ordnance Survey national grid system (aka the British National Grid) (OS 2020). This database is curated by expert from the BBS and was most recently used to published distribution maps for each species (Blockeel et al. 2014). It also forms the main source of data available for this organism group and displayed by the NBN Atlas, UK’s largest repository of publicly available biodiversity data (NBN Trust 2024).

**Figure 1:**
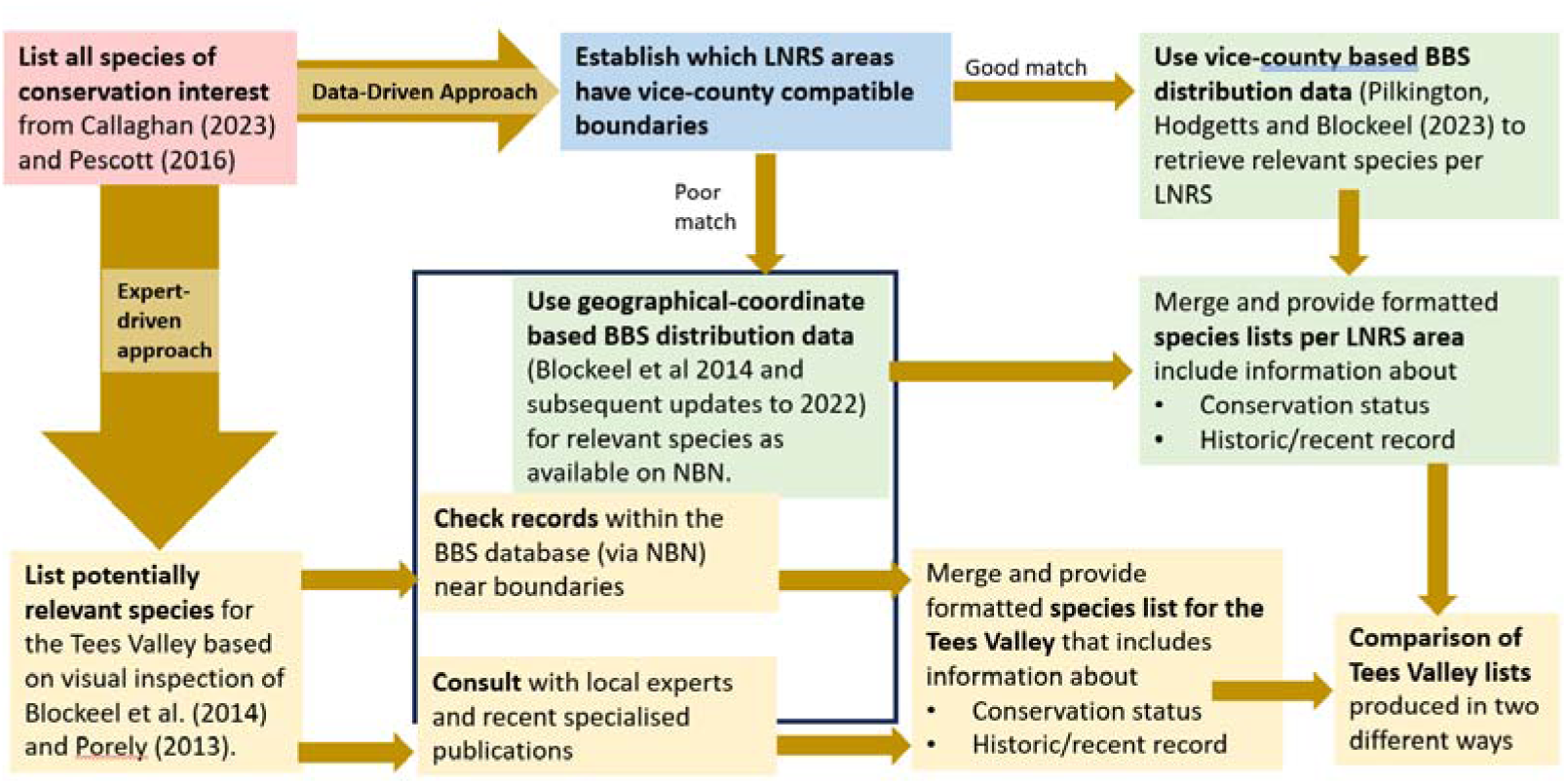
Conceptual flowchart of the work presented. In pink, step relating to the identification of species of conservation interest, in blue the GIS analysis comparing vice-county and LNRS-area boundaries, in green production of LNRS area-specific list of bryophytes and in yellow the steps involved to compare the work with an expertly-driven list for the Tees Valley. The black square includes step using the BBS geographical-coordinate-based database via the NBN Atlas.

**Figure 1.**
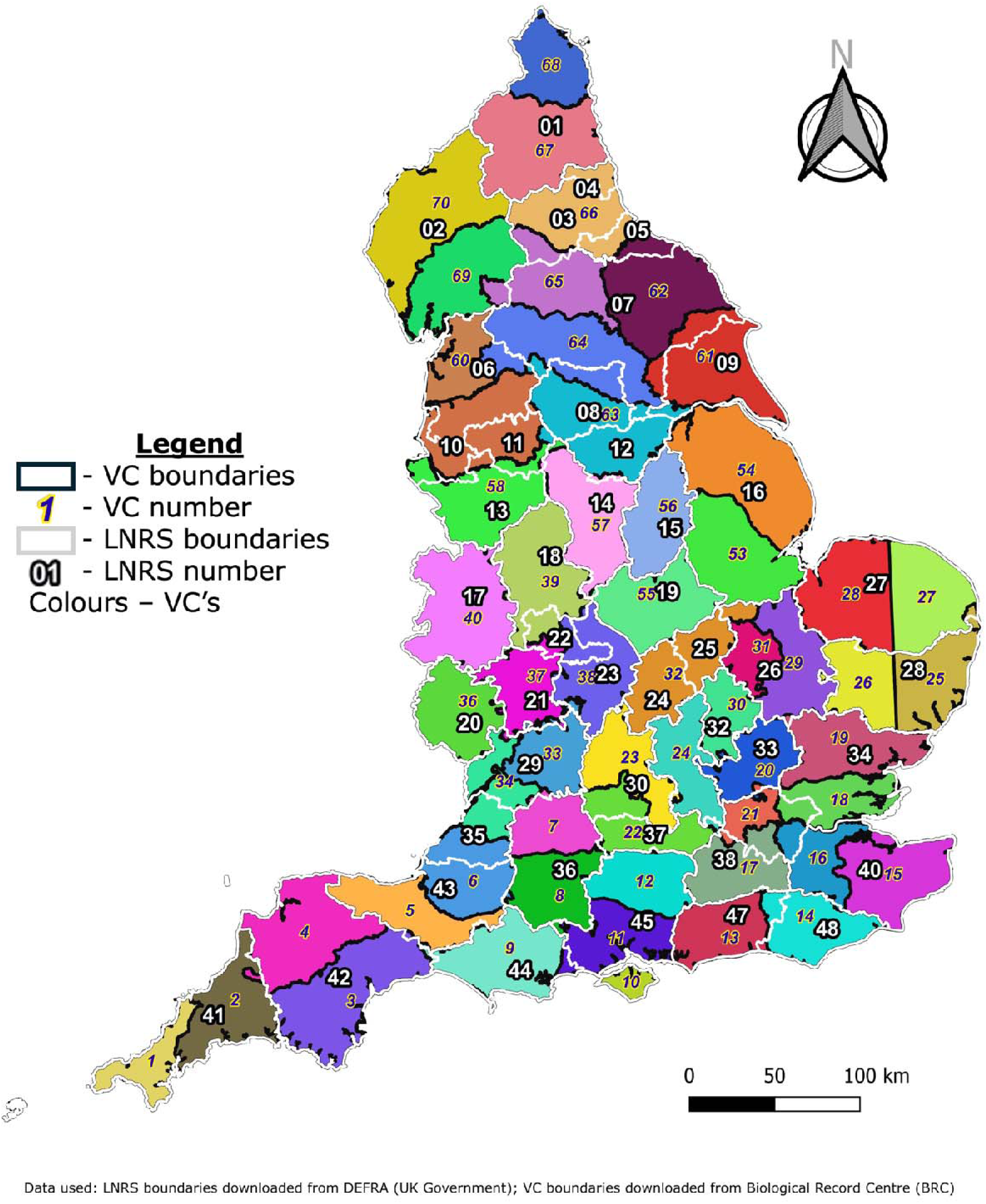
LNRS boundaries overlayed onto coloured VC boundaries. It can be seen that alignment occurs in certain places, but discrepancies are common.

**Figure 2.**
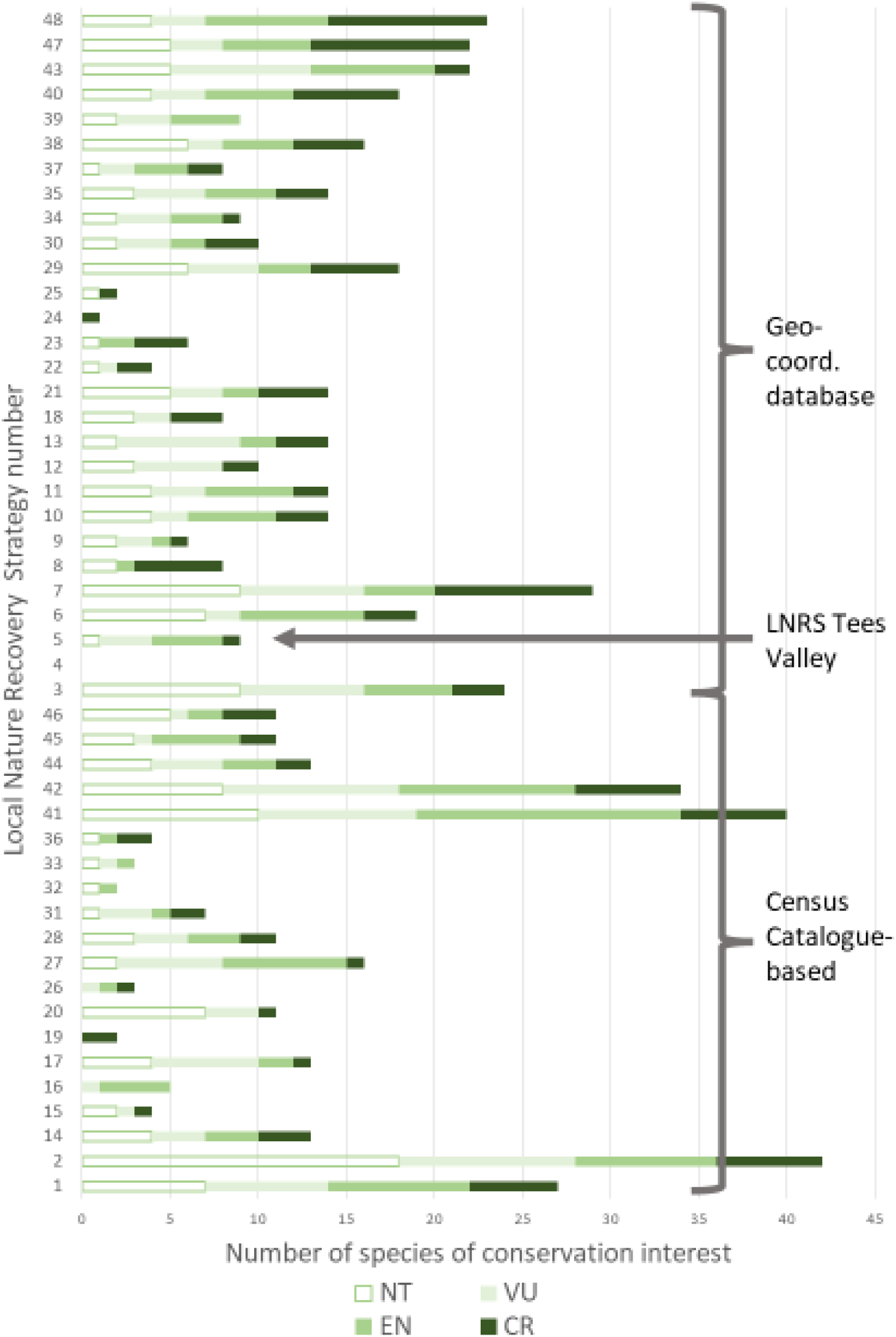
Stacked bar graph displaying the near threatened and threatened species count for each LNRS.

There are four main sections to the methodology used: identifying species of conservation interests, boundary analysis, species list creation and, fourthly, comparison in the Tees Valley. Spatial data visualisation and analysis was conducted using Quantum Geographical Information System (QGIS) Desktop version 3.34.7 ‘Prizren’ (QGIS 2024). For further analysis and data management, Microsoft Excel (Microsoft 2024) and R version 4.4.1 (R Core Team 2024) were used.

### Identifying species of conservation interest

The taxonomic concept adopted was that used in the most recent IUCN Red List of the bryophytes of Britain (Callaghan 2023). Species of conservation interest are defined here as species with status, in Callaghan (2023), identified as either: extinct (EX), extinct in the wild (EW), critically endangered (CR), endangered (EN), vulnerable (VU), near threatened (NT), or data deficient (DD); or listed in Pescott (2016) as nationally scarce (NS) or nationally rare (NR). Despite being partly obsolete after the publication of the IUCN red list, rare and scarce species lists remains integral in identifying conservation sites, including Sites of Special Scientific Interest (SSSI) (Preston 2010; Bainbridge et al. 2013; Bosanquet et al. 2018). Finally, all species included in Section 41 of the Natural Environment and Rural Communities Act 2006 (UK Government 2006) were compared with the list of species conservation interest.

### Boundary analyses

Watsonian vice-county boundaries (BRC 2024; GitHub 2024) and Local Nature Recovery Strategy area boundaries (Natural England 2024) were combined using the ‘intersect’ tool in QGIS, creating GIS features that belonged to only one VC and only one LNRS. For each of these new features (thereafter VC-LNRS features), the surface area (in m2) was calculated and exported. This enabled calculations of the percentage of each VC or LNRS area, that each VC-LNRS feature occupies. Based on these data we devised three thresholds - 95%, 90% and 80% - to determine whether the CC species lists could be considered accurately representative of LNRS areas. A LNRS area qualified at the 95% threshold if it passed two criteria. Firstly, only VC-LNRS features that comprise 95% of the surface area of a VC were considered. Secondly, these features were then used to check which LNRS areas was covered at least to 95% of its total surface area using the relevant VC-LNRS feature(s) selected with the first criteria. The same process was used to determine which LNRS area can be covered by vice counties using a 90% or 80% threshold.

### Creation of data-driven species lists

Using the 90% threshold as an example, lists of species were produced for each LNRS area, using either the CC (>90%) or the BBS database (<90%). Using the CC, the lists are simply a subset of the listed species for a given VC, when they qualify as being of conservation interest. Using the BBS database required further analysis. Observation records from England and for all relevant species were downloaded from the National Biodiversity Network Atlas (NBN Trust 2024). The dataset available includes the entirety of the BBS database as developed for the publication of Blockeel et al. (2014), with additional records made available by the BBS up to the year 2022. To limit uncertainty, records that did not contain at least a 10km grid references were filtered out. The remaining records were attributed to the geographically relevant LNRS areas at the 1km square scale whenever possible, using the 1km square layer of the Ordinance Survey’s (OS) British National Grids, available through the OS GitHub repository (OS 2021). When a record’s OS British National Grid coordinates overlapped more than one LNRS area, this was tagged as not unique (to one specific LNRS area). This means that there was some un-certainty about which LNRS area the record should be attributed to and as a cautionary measure the record was attribute to more than one LNRS area.

### Comparison with expert-driven list in the Tees Valley

An additional approach was used in the Tees Valley LNRS area. The historical and recent presence of all relevant taxa were checked using Blockeel et al. (2014), Porley (2013) and the underpinning data from the BBS database as accessible through the NBN Atlas. Local experts were also contacted for any additional species of interest that may have been observed between 2014 and 2024, or any additional species of conservation interest not highlighted with this process. The resulting list was compared with that obtained with the other data-driven approach for the Tees Valley.

## Results

### Species of conservation interest

The merging of the red list (RL) Callaghan (2023), the nationally rare and scarce (NR/NS) classification (Pescott 2016) and relevant to English vice-counties in the CC (Pilkington et al. 2023) returned a total of 373 species of conservation interest found within England (Appendix 1). Of these, 34 – or 9.1% - are categorised as CR/CR(PE) against Red List criteria; The other two main RL categories, EN and VU, have 36 (9.7%) and 32 species (8.6%) respectively. Under RL criteria over a quarter – 27.3% - of bryophyte species present in England are considered under threat of extinction. All but one species, *Bruchia vogesiaca* Nestl. ex Schwägr., had occurrence data available from the NBN Atlas. That species was spotted in Britain for the first time in 2006 (Holyoak 2007) and appears to be very restricted in distribution or even extinct (Callaghan 2016). With all datasets merged, the total number of observations with 1km or 10km grid references was 30,850, with 343 species (92% of England’s total) included. All species of principal importance under Section 41 (UK Government 2006), were included in our list of species of conservation interest.

### Boundary analysis

Intersecting the two boundary layers succeeded in creating small features, 289 in total, each solely belonging to one individual VC and one individual LNRS area (provided as a GIS layer in Appendix 2). Thirty features had calculated areas under 1 ha, suggesting slight discrepancies e.g. in coastlines digitisation, but this did not constitute evidence of major discrepancies between the two layers. Of the 48 LNRS areas, only the Isle of Man was a 100% match with its vice-county. Against the three thresholds, 17 LNRS areas - 35% - are at or above 95% covered, 20 areas – 42% - meet 90%, and 28 areas – 58% - are an 80% match (Table 1). Due to the small difference between 95% and 90%, it was decided that 90% offered maximum coverage without sacrificing accuracy. The LNRS areas above the 90% threshold are, as expected, mostly along the coasts and the borders with Wales and Scotland (Figure 1), where boundaries are less likely to change over time. Excluded areas with the 90% threshold appearing mostly around large population centres, such as London and Yorkshire. Ten of the twenty LNRS included, i.e. 50%, are made up of two qualifying VC’s.

**Table 1:**
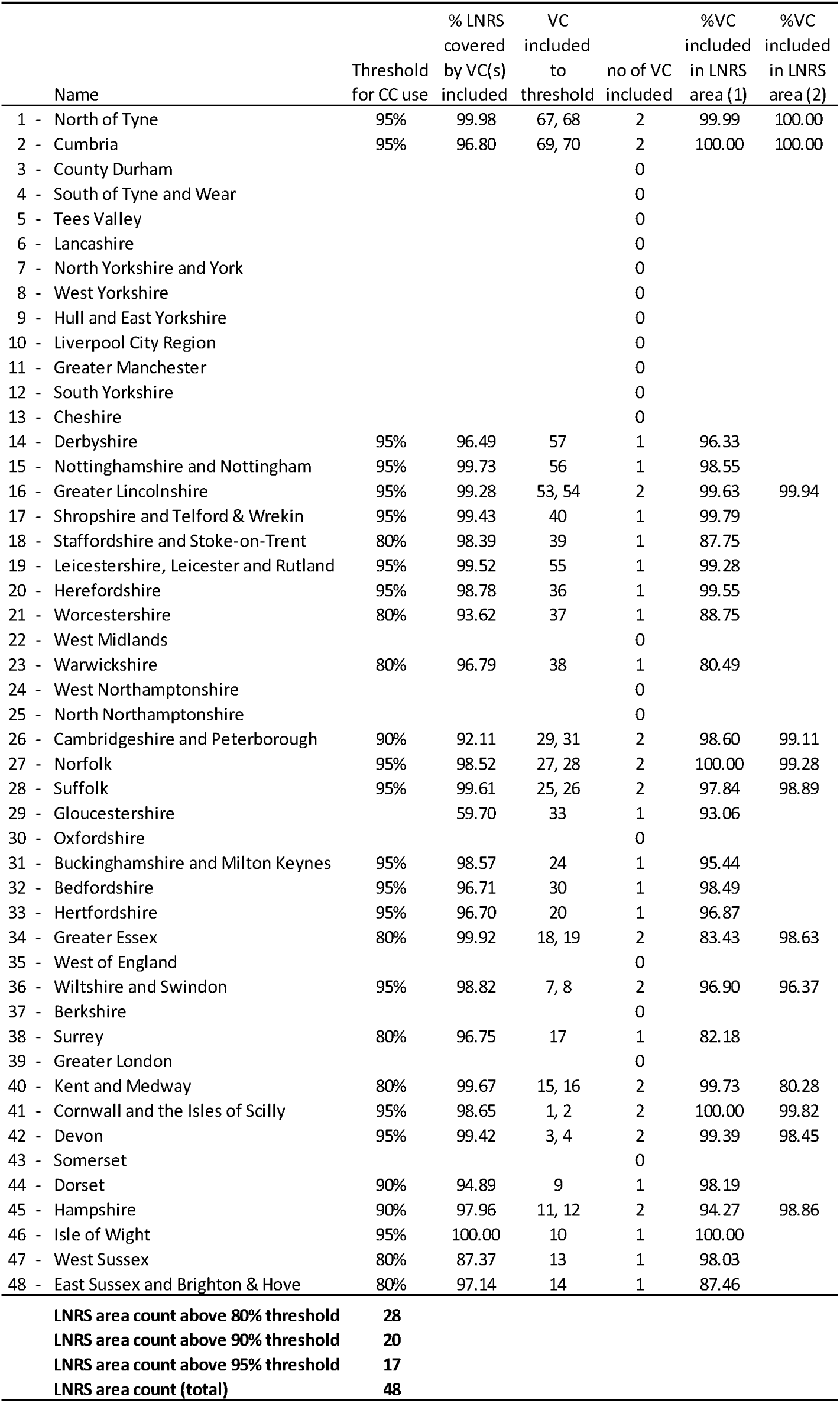
List of LNRS area indicating whether the vice-county based Census Catalogue (CC) could be used, at three threshold of match, 95, 90 and 80%. (See methods for full explanation of inclusion criteria)

### Creation of data-driven species lists

For the 20 LNRS areas compatible with the CC, there was a range of 163 species between the largest and lowest total numbers: LNRS Cumbria, a combination of VC Westmorland and VC Cumberland, having 186 species present and LNRS Bedfordshire, matching VC Bedfordshire, having 23 species. LNRS Cumbria, LNRS Cornwall and the Isles of Scilly, and LNRS Devon all have 6 CR species; LNRS Greater Lincolnshire, LNRS Bedfordshire, and LNRS Hertfordshire all having zero. For total number of species deemed to be at threat of extinction, i.e. those categories as CR, EN, or VU, LNRS Cornwall and the Isles of Scilly has most with 30; with 1, LNRS Bedfordshire had the lowest. Across all LNRS areas in this group, 9.6 was the mean number of at-risk species, with a large standard deviation of 8.7 (Table 2). For NR/NS species, LNRS Cornwall and the Isles of Scilly again had most NR with 45, and LNRS Cumbria had most NS with 153, and the most combined with 181 of 186 (97.3%) species present categorised under this system. The full break down per LNRS area is given in Appendix 3.Twenty-eight LNRS areas did not meet the 90% threshold (Table 3).

**Table 2:** average number of taxa selected within the LNRS area lists and their threat and rarity.

| Species status | Vice-county-based |  | Coordinate-based |  | All areas |  |
| --- | --- | --- | --- | --- | --- | --- |
|  | mean | std dev | mean | std dev | mean | std dev |
| IUCN criteria threat |  |  |  |  |  |  |
| CR | 2.3 | 2.0 | 3.2 | 2.5 | 2.8 | 2.3 |
| EN | 3.8 | 4.0 | 3.0 | 2.3 | 3.3 | 3.1 |
| VU | 3.5 | 3.4 | 2.9 | 2.2 | 3.2 | 2.7 |
| CR, EN or VU | 9.6 | 8.7 | 9.2 | 5.3 | 9.3 | 6.9 |
| NT | 4.1 | 4.4 | 3.4 | 2.5 | 3.6 | 3.4 |
| LC | 52.6 | 27.2 | 47.5 | 22.4 | 49.6 | 24.4 |
| DD | 3.2 | 2.8 | 2.8 | 1.5 | 3.0 | 2.1 |
| NA | 1.2 | 1.3 | 0.3 | 0.8 | 0.6 | 1.1 |
| Nationally rare (NR) and scarce (NS) |  |  |  |  |  |  |
| NR | 13.2 | 10.8 | 10.5 | 6.1 | 11.6 | 8.4 |
| NS | 55.8 | 31.3 | 50.6 | 24.1 | 52.7 | 27.1 |
| Total | 70.5 | 41.0 | 63.3 | 30.5 | 66.3 | 35.1 |

**Table 3.** Species of conservation interest identified as relevant to the Tees Valley LNRS area.

| Species | IUCN / NS / NR | Historic / Recent | Inclusion using data-driven approach | Inclusion using expert driven approach |
| --- | --- | --- | --- | --- |
| *Hedwigia ciliata* | NS | na | Yes, but erroneous (taxonomy) | No |
| *Homomallium incurvatum* | EN | na | Yes, but erroneous (boundaries) | No |
| *Campylostelium saxicola* | NS | na | Yes, but erroneous (boundaries) | No |
| *Hennediella stanfordensis* | NS | na | Yes, but erroneous (boundaries) | No |
| *Herzogiella seligeri* | NS | na | Yes, but erroneous (boundaries) | No |
| *Platydictya jungermannioides* | NS | na | Yes, but erroneous (boundaries) | No |
| *Tortella inclinata* | NS | na | Yes, but erroneous (boundaries) | No |
| *Conardia compacta* | NT | na | Yes, but erroneous (boundaries) | No |
| *Orthothecium rufescens* | NS | Historic only | Yes | Yes |
| *Bryum uliginosum* | CR(PE) | Historic only | Yes | Yes |
| *Drepanocladus sendtneri* | DD(NS) | Recent | Yes | Yes |
| *Bryum calophyllum* | EN | Historic only | Yes | Yes |
| *Bryum marratii* | EN | Historic only | Yes | Yes |
| *Bryum warneum* | EN | Recent | Yes | Yes |
| *Orthotrichum pallens* | NR | Historic only | Yes | Yes |
| *Aloina brevirostris* | NS | Recent | Yes | Yes |
| *Amblyodon dealbatus* | NS | Historic only | Yes | Yes |
| *Brachythecium salebrosum* | NS | Historic only | Yes | Yes |
| *Discellium nudum* | NS | Recent | Yes | Yes |
| *Distichium inclinatum* | NS | Recent | Yes | Yes |
| *Hygroamblystegium humile* | NS | Historic only | Yes | yes |
| *Nardia geoscyphus* | NS | Historic only | Yes | Yes |
| *Rhynchostegiella curviseta* | NS | Historic only | Yes | Yes |
| *Ricciocarpos natans* | NS | Recent | Yes | Yes |
| *Tortula freibergii* | NS | Recent | Yes | Yes |
| *Weissia squarrosa* | NS | Historic only | Yes | Yes |
| *Bryum knowltonii* | VU | Historic only | Yes | Yes |
| *Petalophyllum ralfsii* | VU | Historic only | Yes | Yes |
| *Tomentypnum nitens* | VU | Historic only | Yes | Yes |
| *Oleolophozia perssonii* | NS | Recent | Yes | No, but missing (recent) |
| *Dicranella crispa* | NR | Historic only | Yes | No, but missing (historical) |
| *Fissidens rufulus* | NS | Historic only | Yes | No, but missing (historical) |
| *Pterygoneurum ovatum* | NS | Historic only | Yes | No, but missing (historical) |
| *Pylaisia polyantha* | NS | Historic only | Yes | No, but missing (historical) |
| *Acaulon muticum* | NS | Historic only | No, but missing (not on NBN) | Yes |
| *Aloina ambigua* | NS | Recent | No, but missing (Baker 2022) | Yes |
| *Encalypta rhaptocarpa* | NS | Recent | No, but missing (Baker 2022) | Yes |
| *Moerckia flotoviana* | NS | Recent | No, but missing | yes |
| *Blepharostoma trichophyllum* |  | Recent | No, but not expected | Yes |
| *Encalypta vulgaris* |  | Recent | No, but not expected | Yes |
| *Entosthodon obtusus* |  | Recent | No, but not expected | Yes |
| *Kurzia pauciflora* |  | Recent | No, but not expected | yes |
| *Scorpidium scorpioides* |  | Recent | No, but not expected | Yes |
| *Sphagnum medium* |  | Recent | No, but not expected | Yes |
| *Weissia longifolia* |  | Recent | No, but not expected | Yes |

Across these, the created species lists suggest there is a range of 137 in total number of species, from 8 present in LNRS South of Tyne and Wear to 145 in LNRS North Yorkshire and York. The highest number of CR species present is 9, found across LNRS North Yorkshire and York, LNRS West Sussex, and LNRS East Sussex and Brighton and Hove; two LNRS, South Tyne and Wear and Greater London, have zero. For total number of species deemed at risk of extinction, LNRS North Yorkshire and York is highest with 20 and LNRS South Tyne and Wear has zero. The mean number of at-risk species is 9.2, with a standard deviation of 5.2. Most species classified as both NR and NS are found in North Yorkshire and York, with 26 and 115 respectively. The full break down per LNRS area is given in Appendix 3.

### Comparison with expert-driven list in the Tees Valley

Some 45 species of conservation interest were selected for the LNRS Tees Valley when combining the results of the data-driven and expert driven methods. Of those 45, 21 had been selected using both methods. Some 8 species had been erroneously included into the data-driven list because of method inaccuracies at boundaries when working at 1km scale. Some 7 species has been excluded from the data-driven list because they did not met the inclusion criteria (IUCN or NR/NS). Some 4 species were excluded from the data-driven list because their presence was not included in the BBS geographical coordinate-based database as available via the NBN Atlas at the time of analysis and, finally, some 5 species were not included by the expert-driven assessment due to gross errors. The detail of the species concerned, and their status is presented in table 3.

## Discussion

This study describes a methodology capable of generating relevant bryophyte species lists for each of the forty-eight LNRS areas in England. Our first key objective was to list species of conservation interest in England. We found that under RL criteria over a quarter – 27.3%, 102 species - of bryophyte species present in England are considered under threat of extinction. Adding to this list species previously assessed as nationally rare or scarce, we identified a total of 373 species of conservation interest. This list of species of conservation interest covered all species that are listed in the Section 41 (UK Government 2008). Our second key objective was to ascertain which LNRS areas matched the current recording schemes based on Watsonian vice-counties. Against the three thresholds of 95%, 90% and 80% match between the vice-county boundaries and the 48 LNRS area, 17 LNRS areas, 20 areas and 28 areas, respectively, could benefit from the CC’s database. The 90% threshold was chosen for further analyses. Our third key objective was to produce species lists for each LNRS area and using the most relevant distributional database. We found that LNRS areas had on average 9.3 species threatened of extinction under IUCN criteria and a further 3.6 species on average tagged as near threatened. There was also an average of 3.0 species with a data deficient status and 0.3 species that had not been assessed by Callaghan (2023). There was a further 49.6 species per LNRS area on average that were of least concerned under IUCN criteria but considered as either nationally rare or nationally scarce. Finally, our fourth key objective was to compare the results obtained with the BBS geographical coordinate-based database with those obtained using expert knowledge in the Tees Valley LNRS. The comparison revealed significant differences, with only 21 species selected by both approaches but a total of 45 species listed altogether. The differences could be explained mostly by simple method used at boundaries using the 1-km analysis scale and by additional species being identified by expert knowledge, which did not have any formal conservation status.

### Identifying species of conservation interest

Bosanquet et al. (2018) listed 474 bryophyte species relevant when selecting Sites of Special Scientific Interest, which contrasts with the 373 species we identified as relevant to LNRSs. However, the geographical scope of their assessment is the whole of Britain, including Scotland and Wales, and the uplands of these countries are known to be bryological diversity hotspots. Thus, the additional species not listed in our work can be assumed not to occur on England. Another consideration is that we were able to use the latest IUCN-criteria red list (Callaghan, 2023), which had not been fully updated and published in 2018. In contrast, relying on Pescott (2016) for nationally rare and scarce species means that a small number of species did not benefit from the latest distributional knowledge in our assessment. Importantly, our list included all species listed in the Schedule 41 (UK Government 2006), bringing additional insurance that our approach covered appropriately species of conservation value.

### Boundary analyses

It is remarkable to think that after nearly 200 years the Watsonian vice-counties and their use in biological recording are relevant to nature recovery in 20 out of 48 areas of England. Our work also highlights that the coordinate-based database of the BBS is more nimble and adaptable for extracting data when new policies arise. This agility, however, comes at a cost, as additional bioinformatics skills are required to extract the data in a suitable format for a new policy or scheme. Spatially-informed bioinformatics tools have been highlighted for over a decade as capital when aiming to improve our knowledge of biodiversity distribution, and when managing global change (e.g. Jetz et al. 2012), because there can be discrepancy of scales between distributional knowledge and information needed on the ground for conservation action. In addition, resources for planning nature recovery are scarce and the specialised skills to integrate distributional knowledge into strategy documents may not always be systematically available for each local area. It is also to be noted that one disadvantage of coordinate-based data in our exercise was that some records did not have coordinates at 10km scale or lower. Some 29 species (about 8%) were lost following the grid reference filtering processes, because none of their occurrence records provided 1km or 10km grid references. Lacking grid references is likely to concern species that have not been found recently and whose past presence is only attested by herbarium specimens without precise location entered directly into the BBS database. However, expert knowledge and specialised publication such as Porley (2013) could be used to fill some of these gaps. Poor location recording is a known issue, and more standard data collection of geographical locations has been proposed by Callaghan (2022). For species of conservation interest, it is recommended to record to accuracy of 10m and 1m squares or individual trees (Callaghan 2023). Using this granular data would allow more in-depth spatial analysis to be conducted, and eventually modelling that considers elevation, slope, surrounding vegetation type, soil quality, proximity to water sources, microclimate, and other ecologically important variables (Tenedorio and Rocha 2018). Armed with a full array of variables, it would be possible to perform desk-based assessments not only of species and habitat types, but ecosystem services, another vital aspect of considering areas for protection or conservation (Magdalena et al. 2022; DEFRA 2023). For LNRS, this would enable, for example, narrowing down specific habitats to prioritise.

### Creation of data-driven species lists

The results of this project may be of interest to a wide array of people and organisations. Bryophytes are not only often overlooked (Boch et al. 2013), but also noted as a taxonomic group suffering from a general lack of knowledge and/or interest (Slate et al. 2024) and even being poorly understood by those engaged in studying them (Eldridge et al. 2023). With not every local authority employing an in-house professional ecologist (ALGE 2024), or having access to a wide array of experts operating locally, possessing reference material regarding priority biodiversity data tailored to individual areas, could be a valuable resource. The results highlight a high number of relevant species when applying the recommended criteria (Natural England 2023), even when restricting the searches to species on the brink of extinction (i.e. IUCN categories CR, EN and VU). However, there was a large amount of variability between areas (see Table 3).

The successful creation of bespoke bryophyte species lists for each LNRS points to the viability of the designed methodology. This is not just useful for bryophytes, but directly transferable to occurrences of other species recorded against VC boundaries. Two examples are fungi, such as Rusts (Woods et al. 2015), and insects, such as Soldier flies and Allies (Dipterists 2024). Much like non-vascular plants, certain insect groups receive less attention than the most charismatic species. Big, beautiful and colourful, butterflies and bees tend to dominate insect conservation conversations to the detriment of flies, despite their vitally important ecological roles as detritivores and predators, and as a food source (Dipterists 2024). Ultimately, the more relevant species brought to attention of environmental managers at local scales, the more effective LNRS are likely to be at the nation-scale.

### Comparison with expert-driven list in the Tees Valley

It come with no surprise that expert knowledge delivers more granularity and more up to date knowledge, while the data-driven approach is more reproducible and based on publicly available data. The expert-driven approach practiced a sharper taxonomic assessment and enabled checks at finer geographical scales near the LNRS area. For example, the relevant record of *Hedwigia ciliata* for the Tees Valley referred to a broader taxonomic concept for this species (*Hedwigia ciliata sl*) that has now been abandoned (Hedenas 1994). As a results, in all likelihood, the species present in the North York Moors and within the Tees Valley LNRS is *Hedwigia stellata* Hedenaes (Crundwell 1995), a species that is not a conservation priority. Similar errors will be found throughout the work presented here for this taxa and other taxa that experienced complex taxonomic changes in recent decades. However, with an improved algorithm and better attention to taxonomic detail, many of these errors could be eliminated from the data-driven method in the future. The situation is similar for species that the expert-driven approach excluded based on finer scales than the 1km grid square we used with the data-driven approach. For example, expert opinion enabled the exclusion of *Conardia compacta* (Müll.Hal.) H.Rob., which is reported for a 1km grid square straddling the Tees Valley LNRS boundaries but from the toponym “Castle Eden Dene” which is fully outside the boundaries. In many instances, rare plants such as those selected here as being of conservation interest, and especially those reported in recent decades, will be associated with coordinates at a finer scale than 1km grid squares and therefore the data-driven methodology can be improved in the future. One aspect that would be difficult to improve for the data-driven appraoch, is the reference to very recent data and discoveries that have not entered the public domain, e.g. via the NBN Atlas. In our case, a consultancy report (Anonymous, 2010), as well as field work conducted by the British Bryological Society in the Tees Valley (Baker 2023) both highlighted a few species of importance for the LNRS, such as recent records of *Bryum warneum* (Röhl.) Brid, a species categorised as Endangered in Britain by Callaghan (2023) and as Vulnerable at the European level (Hodgett et al. 2019). Although greatly improved by the advent of bio-informatics, there will always remain a time lag before natural history sightings and publication, typically verified, compiled and uploaded by volunteer citizen scientists. Despite this drawback, the data-driven approach proved more reliable in five specific instances, when highlighting gross errors in the form of omission in the expert-driven assessment (Table 3). Finally, some 8 taxa were included by expert opinion but do not have any formal conservation status. *Enthostodon obtusus* (Hedw.) Lindb., for example, is absent or declining from most parts of England and its presence in the North York Moors parts of the Tees Valley LNRS area is of English significance (Blockeel et al. 2014).

However, at the British scale, the species is less rare being widely reported from Wales and Western Scotland. This highlights the discrepancy in geographical scope between the LNRS being relevant to England only, while the publications used to assess the conservation value are based on Britain as a whole (Callaghan 2023; Pescott 2016), albeit without the UK’s Northern Ireland nation. However, using a British red list brings additional guaranties that local scale nature recovery efforts through the LNRS are also of broader international relevance, Britain being one of the bryophyte biodiversity hotspots in Europe (Hodgetts, et al 2020). On the other hand, ignoring this discrepancy in geographical scope within the current role may results in ignoring some species of conservation interest within England but that are relatively common in other parts of Britain. This latter risk could be mitigated by making widely available the findings of Porley (2013) and any further updates, where the status of England’s rarest bryophytes was rigorously analysed. Based on the above, it appears that both expert opinion and data-driven approaches have advantages and drawback, suggesting that a combination of both would be beneficial. Specifically, we suggest that the data-driven approach presented here will save considerable amount of time and accuracy to local experts, who inevitably will be led to improve the focus of LNRS in the future.

## Acknowledgments

The authors wish to thank the LNRS Tees Valley species specialist group members, as well as colleagues at Teesside University for discussion and inspiration for this work.

