## Appendix 2 for "Developing an approach to list bryophytes of conservation interests for England’s Local Nature Recovery Strategies": Intersections_LNRS_VC.qmd

```
<!DOCTYPE qgis PUBLIC 'http://mrcc.com/qgis.dtd' 'SYSTEM'>
<qgis version="3.34.7-Prizren">
  <identifier></identifier>
  <parentidentifier></parentidentifier>
  <language></language>
  <type>dataset</type>
  <title></title>
  <abstract></abstract>
  <contact>
    <name></name>
    <organization></organization>
    <position></position>
    <voice></voice>
    <fax></fax>
    <email></email>
    <role></role>
  </contact>
  <links/>
  <dates/>
  <fees></fees>
  <encoding></encoding>
  <crs>
    <spatialrefsys nativeFormat="Wkt">
      <wkt>PROJCRS["OSGB36 / British National Grid",BASEGEOGCRS["OSGB36",DATUM["Ordnance Survey of Great Britain 1936",ELLIPSOID["Airy 1830",6377563.396,299.3249646,LENGTHUNIT["metre",1]]],PRIMEM["Greenwich",0,ANGLEUNIT["degree",0.0174532925199433]],ID["EPSG",4277]],CONVERSION["British National Grid",METHOD["Transverse Mercator",ID["EPSG",9807]],PARAMETER["Latitude of natural origin",49,ANGLEUNIT["degree",0.0174532925199433],ID["EPSG",8801]],PARAMETER["Longitude of natural origin",-2,ANGLEUNIT["degree",0.0174532925199433],ID["EPSG",8802]],PARAMETER["Scale factor at natural origin",0.9996012717,SCALEUNIT["unity",1],ID["EPSG",8805]],PARAMETER["False easting",400000,LENGTHUNIT["metre",1],ID["EPSG",8806]],PARAMETER["False northing",-100000,LENGTHUNIT["metre",1],ID["EPSG",8807]]],CS[Cartesian,2],AXIS["(E)",east,ORDER[1],LENGTHUNIT["metre",1]],AXIS["(N)",north,ORDER[2],LENGTHUNIT["metre",1]],USAGE[SCOPE["Engineering survey, topographic mapping."],AREA["United Kingdom (UK) - offshore to boundary of UKCS within 49°45'N to 61°N and 9°W to 2°E; onshore Great Britain (England, Wales and Scotland). Isle of Man onshore."],BBOX[49.75,-9.01,61.01,2.01]],ID["EPSG",27700]]</wkt>
      <proj4>+proj=tmerc +lat_0=49 +lon_0=-2 +k=0.9996012717 +x_0=400000 +y_0=-100000 +ellps=airy +units=m +no_defs</proj4>
      <srsid>2437</srsid>
      <srid>27700</srid>
      <authid>EPSG:27700</authid>
      <description>OSGB36 / British National Grid</description>
      <projectionacronym>tmerc</projectionacronym>
      <ellipsoidacronym>EPSG:7001</ellipsoidacronym>
      <geographicflag>false</geographicflag>
    </spatialrefsys>
  </crs>
  <extent>
    <spatial maxy="-179769313486231570814527423731704356798070567525844996598917476803157260780028538760589558632766878171540458953514382464234321326889464182768467546703537516986049910576551282076245490090389328944075868508455133942304583236903222948165808559332123348274797826204144723168738177180919299881250404026184124858368" crs="EPSG:27700" maxz="0" maxx="-179769313486231570814527423731704356798070567525844996598917476803157260780028538760589558632766878171540458953514382464234321326889464182768467546703537516986049910576551282076245490090389328944075868508455133942304583236903222948165808559332123348274797826204144723168738177180919299881250404026184124858368" dimensions="2" minx="179769313486231570814527423731704356798070567525844996598917476803157260780028538760589558632766878171540458953514382464234321326889464182768467546703537516986049910576551282076245490090389328944075868508455133942304583236903222948165808559332123348274797826204144723168738177180919299881250404026184124858368" miny="179769313486231570814527423731704356798070567525844996598917476803157260780028538760589558632766878171540458953514382464234321326889464182768467546703537516986049910576551282076245490090389328944075868508455133942304583236903222948165808559332123348274797826204144723168738177180919299881250404026184124858368" minz="0"/>
    <temporal>
      <period>
        <start></start>
        <end></end>
      </period>
    </temporal>
  </extent>
</qgis>
```
